# Perceived stress exacerbates psoriasis in human skin *in vivo*: Insights from a humanized psoriasis mouse model

**DOI:** 10.1101/2024.09.22.614301

**Authors:** Assaf A Zeltzer, Aviad Keren, Samieh Shinnawi, Marta Bertolini, Ralf Paus, Amos Gilhar

## Abstract

**Introduction:** The widely held belief that psychoemotional stress triggers or exacerbates psoriatic skin lesions lacks sufficient scientific evidence. This study investigated this concept using a psoriasis humanized mouse model.

**Methods:** Healthy human skin was grafted onto SCID/beige mice (n=25), and one month later, psoriatic lesions were induced by intradermal injection of autologous, in vitro IL-2- preactivated PBMCs. Following lesion development, topical dexamethasone (DEX) was applied to induce lesion remission. After lesions disappeared, the mice were exposed to either sonic or sham stress for 24 hours.

**Results:** Sonic stress led to the relapse of psoriatic lesions in all human skin xenografts within 14 days. This relapse was associated with significant changes in psoriasis-related skin characteristics: increased epidermal thickness, K16 expression, keratinocyte proliferation, antimicrobial peptide expression (S100A7, hβ2-defensin), and immune activation markers (HLA-DR, ICAM-1, CD1d, MICA-NKG2D). Additionally, epidermal and dermal immune cells (CD3+, CD8+, CD11c+, CD56+, ILC3, c-KIT+ or tryptase+ cells) and psoriasis-associated pro-inflammatory mediators (CXCL10, IL-22, IL-15, IL-17A/F, IFN-γ, and TNFα) were found to be increased. Neurogenic inflammation biomarkers (NGF, NK1-R, and substance P) were also significantly upregulated in stressed mice. Treatment with the FDA-approved neurokinin-1 receptor antagonist, aprepitant, prevented stress-induced psoriatic relapses in 4 out of 5 mice and normalized most inflammatory and neurobiological markers.

**Conclusions:** These findings provide novel, conclusive evidence that perceived stress can trigger psoriatic lesions in human skin xenografts in vivo and highlight the role of substance P-dependent neurogenic inflammation in this process.

---

Even though many affected patients are convinced that ‘stress’ can trigger or exacerbate their psoriasis lesions, it has long been controversially debated if psychoemotional (perceived) stress can indeed do this in human skin, namely by promoting neurogenic skin inflammation (Griffiths et al., 2021; Roger et al., 2022; Marek-Jozefowicz et al., 2022, Snast et al., 2018; S1-S3).

To address this open question, we have further developed our established humanized psoriasis mouse model (Keren et al., 2018), which mimics human psoriasis as closely as possible *in vivo* (Gilhar et al., 2023), by administering a well-defined perceived stressor, i.e. sound-induced (sonic) stress (Keren et al., 2023; S4). Human skin xenotransplants not only retain core human immune cell populations after transplantation onto SCID/beige 4 weeks post-transplantation and rapidly get re- innervated by sensory mouse nerve fibers within this window, but also show neurogenic skin inflammation after exposing the host mice to sonic stress *in vivo* (Keren et al., 2023; S5).

In this pilot study, we have xenotransplanted healthy human abdominal skin, twenty-five, 1□ cm^2^/0.4 □ mm thick skin fragments (one per mouse) derived from two female volunteers, aged 39 and 48 years, who underwent cosmetic plastic surgery, onto 8 weeks old 25 female SCID/beige mice, under appropriate institutional review board approval and animal license. Post-transplantation, the mice were divided into five groups (see **Figure 1a** for details) to assess psoriasis lesions within the xenotransplants after these had first been experimentally induced by autologous, IL-2- activated PBMCs and then clinically resolved by topical dexamethasone treatment, as described (S6). Four days later, test mice were exposed to sonic stress for 24 hours (Conrad Electronics, Berlin, Germany; 300 Hz, 75 dB) at 15-second intervals as described (S7) in the presence/absence of the neurokinin-1 receptor antagonist, aprepitant (S8). Aprepitant was chosen to block the key stress- and neurogenic inflammation-associated neuropeptide, substance P (SP) (Hegron et al., 2023;S2,S7- S9).

**Figure 1.**
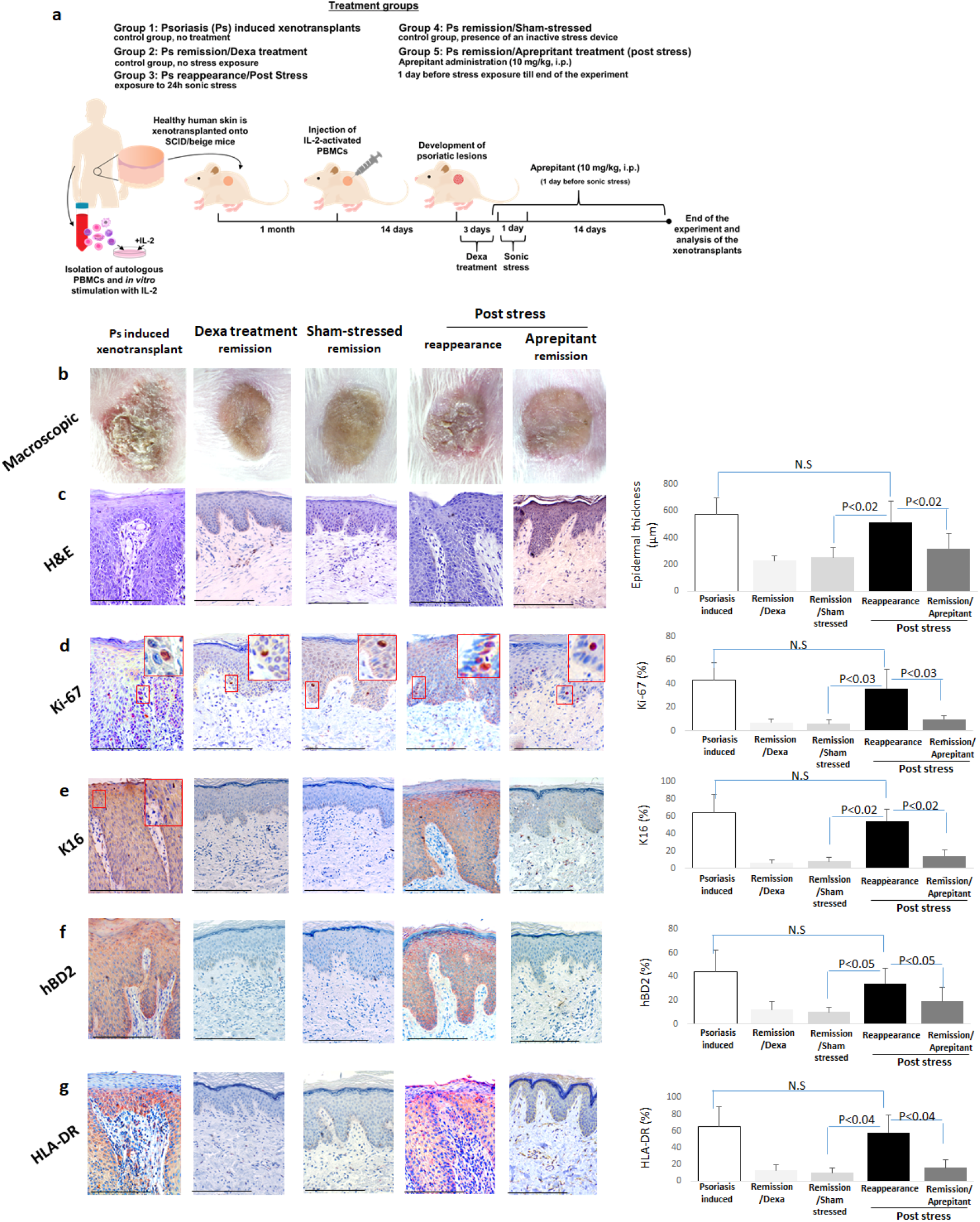
Development of psoriasis lesions in human skin xenotransplants in vivo under sonic stress and aprepitant. Schematic overview of the experimental design. Mice were divided as follows: Group 1 was the control sham treatment, Group 2 lived in a stress-free environment, Group 3 mice were exposed to sonic stress for 24 hours (Keren et al., 2023), Group 4 was exposed to an inactive stress device (sham stress-treatment), and Group 5 received aprepitant prior to sonic stress exposure. Following a period remission after Dexa treatment, the xenografts were analyzed 14 days post-stress as previously described (Keren et al., 2023). Macroscopic and (**c**) histological feature of psoriasis and remission in the psoriasis-induced xenotransplants. Epidermal protein expression of (**d**) Ki-67, (**e**) K16, (**f**) hBD2, and (**g**) HLA-DR were determined by IHC in 25 skin fragments, each derived from 2 independent human donors. For each skin fragment, three non-consecutive skin sections were analyzed. Within each section, four microscopic fields were evaluated at X20 magnification. The pooled means of these measurements were compared using a Student’s t-test, with p-values less than 0.05 being considered statistically significant. In the x-axis of the graphs, ‘remission’ is defined by the reduction or absence of lesions as determined by pathological assessments, indicating decreased disease activity. ‘Reappearance’ refers to the resurgence of disease features through these assessments after a period of remission, signaling an increase in disease activity. Scale bars represent 50 µm. Abbreviations: Dexa, Dexamethasone; IHC, Immunohistochemistry; K16, Keratin 16; hBD2, Human Beta-Defensin 2

This demonstrated significant exacerbation of psoriasis lesions in the xenotransplants following exposure to perceived stress, compared to sham-stressed controls, as evidenced by the elicitation of the characteristic macroscopic and histological features of psoriasis, such as erythema, scaling, increased skin thickness (**Figure 1b**), parakeratosis with absence of granular layer, hyperkeratosis, acanthosis, elongated rete ridges, suprapapillary epidermal thinning, Munro’s microabcesses, dilated tortuous blood vessels, and a pronounced lymphocytic infiltrate (**Figure 1c**).

Quantitative immunohistomorphometry confirmed the expected epidermal hyperproliferation (Ki-67), and significant epidermal overexpression of cytokeratin K16, hβ2-defensin (**Figures 1d-f**), S100A7/psoriasin (**Supplementary Figure 1a**), HLA-DR (**Figure 1g**), ICAM-1, CD1d, and CXCL10 (**Supplementary Figure 1b- d**), in human skin xenotransplants on mice exposed to perceived stress. HLA-DR, ICAM-1, CD1d, and CXCL10 were also overexpressed in the papillary dermis (**Supplementary Figure 1b–d**). Moreover, the number of CD8+ (**Figure 2a**), CD3+ T cells (**Supplementary Figure 1e**), and CD11c+ dendritic cells (**Supplementary Figure 1f**) was significantly increased in the xenotransplants of stressed mice.

**Figure 2.**
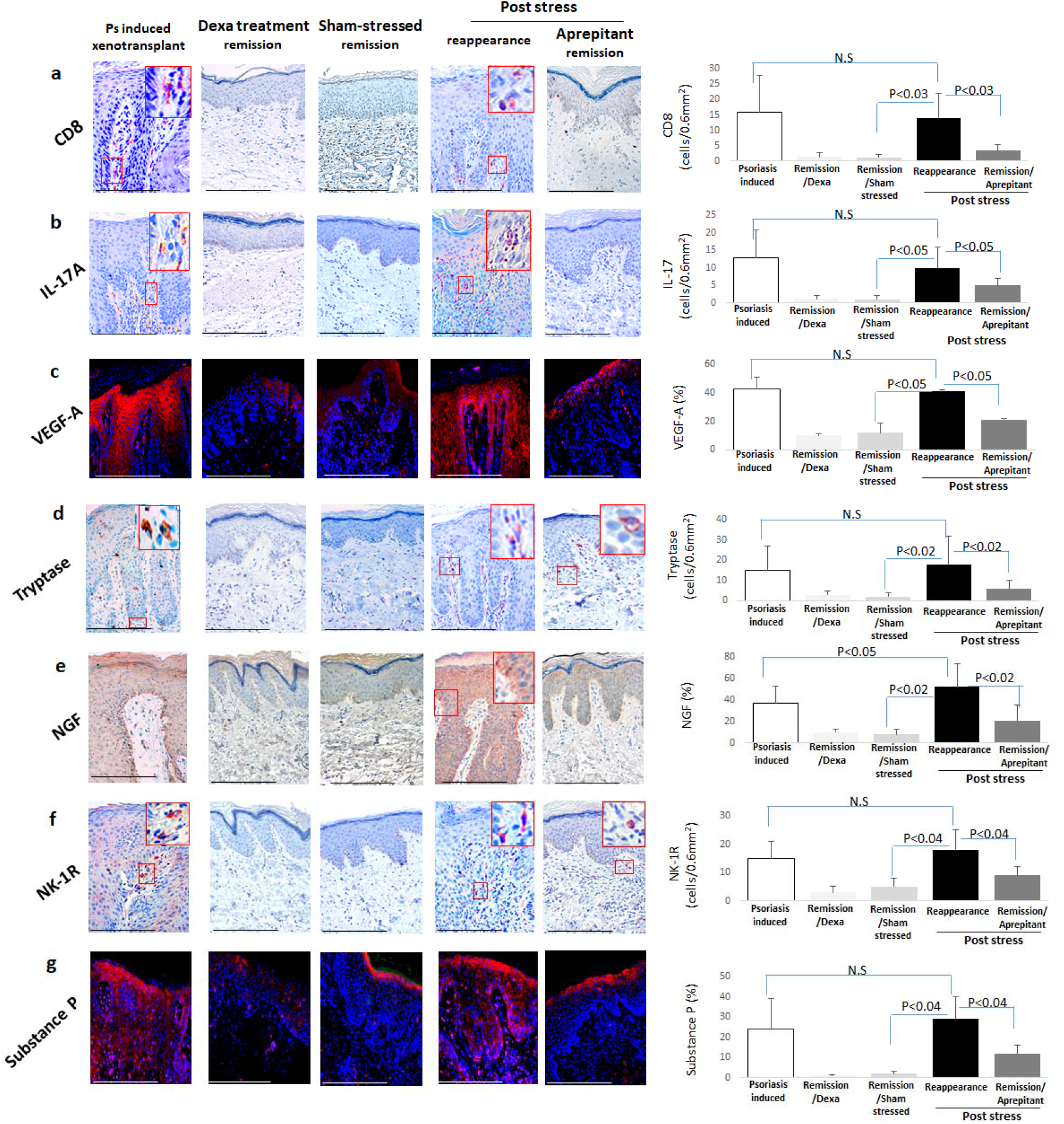
Immunohistochemical profiling in human skin xenotransplants in vivo under sonic stress and aprepitant. This analysis was performed on xenotransplants derived from normal human skin induced to develop psoriatic characteristics, specifically in response to sonic stress and subsequent aprepitant treatment. Markers quantified include (**a**) CD8+ T cells, (**b**) IL-17A, (**c**) VEGF-A, (**d**) Tryptase, (**e**) NGF, (**f**) NK-1R, and (**g**) SP. These were assessed on 25 skin fragments, with three non-consecutive skin sections in total, taken from two independent human skin donors. Within each section, four distinct areas were evaluated at X20 magnification. The data are presented as pooled means, with statistical significance determined by Student’s t-test with p values of less than 0.05 considered statistically significant. The quantitative data reflect the expression levels of dermal CD8+ cells, IL-17A, Tryptase, and NK-1R, as well as epidermal VEGF-A, NGF, and SP. Scale bars: 50 µm. Il, interleukin; NGF, nerve growth factor; Neurokinin-1 Receptor, NK-1R; SP, Substance P; VEGF-A, Vascular Growth Factor-A.

The number of innate immune cells, namely of ILC3 (Keren et al., 2018; S10) and CD56+ cells (**Supplementary Figure 2a and b**), was also significantly increased (S11; S12). Finally, the number of immune cells positive for key cytokines in psoriasis pathogenesis, i.e. IL-22 (**Supplementary Figure 2c**), IL-17A (**Figure 2b**), IL-15, IFN-γ, or TNF-α (**Supplementary Figure 2d–f**), was significantly increased.

Thus, lesions in the human skin xenotransplants of sound-stressed mice showed a wide range of characteristic abnormalities found in psoriasis patients (Gilhar et al., 2023, S13, S14, Gharaee-Kermani et al., 2022).

Given that vascular endothelial growth factor-A (VEGF-A) secretion promotes inflammatory angiogenesis and blood vessel permeability in psoriasis (Luengas- Martinez et al., 2022, S15), it is noteworthy that perceived stress also increased VEGF-A (**Figure 2c**) and matrix metalloproteinase 1 (**Supplementary Figure 2g**) protein expression in the xenotransplants. Thus, perceived stress may facilitate inflammatory cell extravasation, and pathological angiogenesis in psoriasis (S16).

Compared to xenotransplants on sham-stressed mice, tryptase (**Figure 2d**) and c-KIT protein expression (**Supplementary Figure 2h**), pro-nerve growth factor (NGF-β) and SP in the epidermis and neurokinin 1 receptor (NK-1R) in the dermis (**Figure 2e– g**), all of which are up-regulated during neurogenic skin inflammation (Marek-Jozefowicz et al., 2022; S17 -S22), were also significantly increased in xenotransplants on sound-stressed mice. In contrast, administering aprepitant (10 mg/kg, daily) intraperitoneally (S23) to host mice one day before and over 15 days after stress induction, prevented the recurrence of stress-induced psoriasis lesions and significantly reduced psoriasis-associated inflammatory and neurogenic markers (**Figures 1g, Figure 2a,b,d-g, Supplementary Figure 1b-f**; **Supplementary Figure 2a-h, Supplementary Table 1**). This pilot study demonstrates that a) perceived stress can exacerbate psoriatic lesions in adult human skin *in vivo* as a primary trigger factor, supporting circumstantial clinical evidence (Griffiths et al., 2021; Marek- Jozefowicz et al., 2022); b) applying sonic stress in this humanized psoriasis mouse model offers an instructive, novel research tool for preclinically interrogating the insufficiently understood mechanisms that underlie psoriasis exacerbation driven by neurogenic inflammation in human skin *in vivo*, which cannot be reproduced in pure ‘psoriasis’ mouse models (Roger et al., 2022; Gilhar et al. 2023); c) psoriasis exacerbation by perceived stress does not seem to require a distinct genetic predisposition; d) SP/NK-1R-mediated signaling plays a pivotal role in perceived stress-induced relapses of psoriasis, in line with previous research (S2;S13); and e) the FDA-licensed drug, aprepitant, can be repurposed to suppress stress-induced psoriasis relapses in human skin *in vivo*.

Our study is limited by the fact that, owing to the autologous study design, only healthy skin from two female patients, who could provide both skin and PBMCs, was available for transplantation. Thus, study reproduction in a larger *n* of donors and mice, of both genders, in different ethnicities, and under exposure to distinct perceived experimental stressors is required. Also, even though neurogenic skin inflammation was prominently elicited in the human skin xenotransplants *in vivo*, murine nerve fibers and neuromediators are interacting in our model with human skin and its resident immune cell populations. Nevertheless, the current pilot study strongly suggests that ‘stress’-induced psoriasis is quite real - and that pharmacological interventions to counter it can be tested preclinically in this humanized psoriasis model under clinically relevant *in vivo* conditions.

## Abbreviations

PBMCs: peripheral blood mononuclear cells
NGF: nerve growth factor
NK-1R: neurokinin 1 receptor
SP: Substance P
VEGF-A: vascular endothelial growth factor-A
MMP-1: matrix metalloproteinases 1
ILC3: Group 3 innate lymphoid cells.

## Acknowledgements

The authors gratefully acknowledge Prof Petra Arck, University of Hamburg, for kindly donating the sonic stress device used here.

## Data Availability Statement

No complex, website-deposited datasets were generated or analyzed during this study

## Conflict of Interest Statement

The authors state no conflict of interest. For the record, AG performs preclinical psoriasis contract research for competing industry clients, MB is the CEO of Monasterium, while RP is CEO of CUTANEON, a company that develops dermatotherapeutics.

## Author Contribution Statement

The study was conceived and designed by AG and RP, and supervised by AG. Experiments were performed by AK and SS. AZ provided human skin and MB helped with manuscript editing. The manuscript was drafted by AG and RP.

## Notes

### Competing Interest Statement

The authors have declared no competing interest.

