## Supplemental figures for "Perceived stress exacerbates psoriasis in human skin *in vivo*: Insights from a humanized psoriasis mouse model"

**Supplementary Figure Legends**

**Supplementary Figure 1: Quantitative immunohistochemical analysis of psoriatic markers in remission and relapse phases: impact of perceived stress exposure.** Epidermal expression of **(a)** S100A7, **(b)** ICAM,-1, **(c)** CD1d, **(d)** CXCL10 and dermal **(e)** CD3, (**f**) CD11c as determined by IHC performed and quantified on 25 skin fragments, with 3 non-consecutive skin sections in total, taken from two independent human skin donors. Within each section, four distinct areas were evaluated at X20 magnification. The data are presented as pooled means, with statistical significance determined by Student’s t-test with p values of less than 0.05 considered statistically significant. Scale bars: 50 µm.

**Supplementary Figure 2: Immunohistochemical profiling of immune and inflammatory markers in psoriatic induced xenotransplants. (a)** ILC3, **(b)** CD56, **(c)** IL-22, **(d)** IL-15, **(e)** IFN-γ , (**f**) TNF-α, (**g**) MMP-1 and (**h**) c-KIT as determined by IHC performed and quantified on 25 skin fragments, with 3 non-consecutive skin sections in total, taken from two independent human skin donors. Within each section, four distinct areas were evaluated at X20 magnification. The data are presented as pooled means, with statistical significance determined by Student’s t-test with p values of less than 0.05 considered statistically significant. Scale bars: 50 µm. ILC3, Group 3 innate lymphoid cells; TNF-α, Tumor necrosis factor α; MMP-1, matrix metalloproteinase 1.

**Supplementary Table 1: Summary of psoriasis-associated biomarker expression**. The table provides a comprehensive overview of the expression levels of defined psoriatic biomarkers under different treatment and stress conditions in human skin xenotransplants with experimentally induced psoriasis lesions *in vivo* (experimental design: see **Fig. 1a**). Measurements by quantitative immunohistomorphometry are indicated as mean ± standard deviation, and the mean number of cells per 0.66mm² reference area at x20 magnification is listed for selected read-outs.

Top of Form

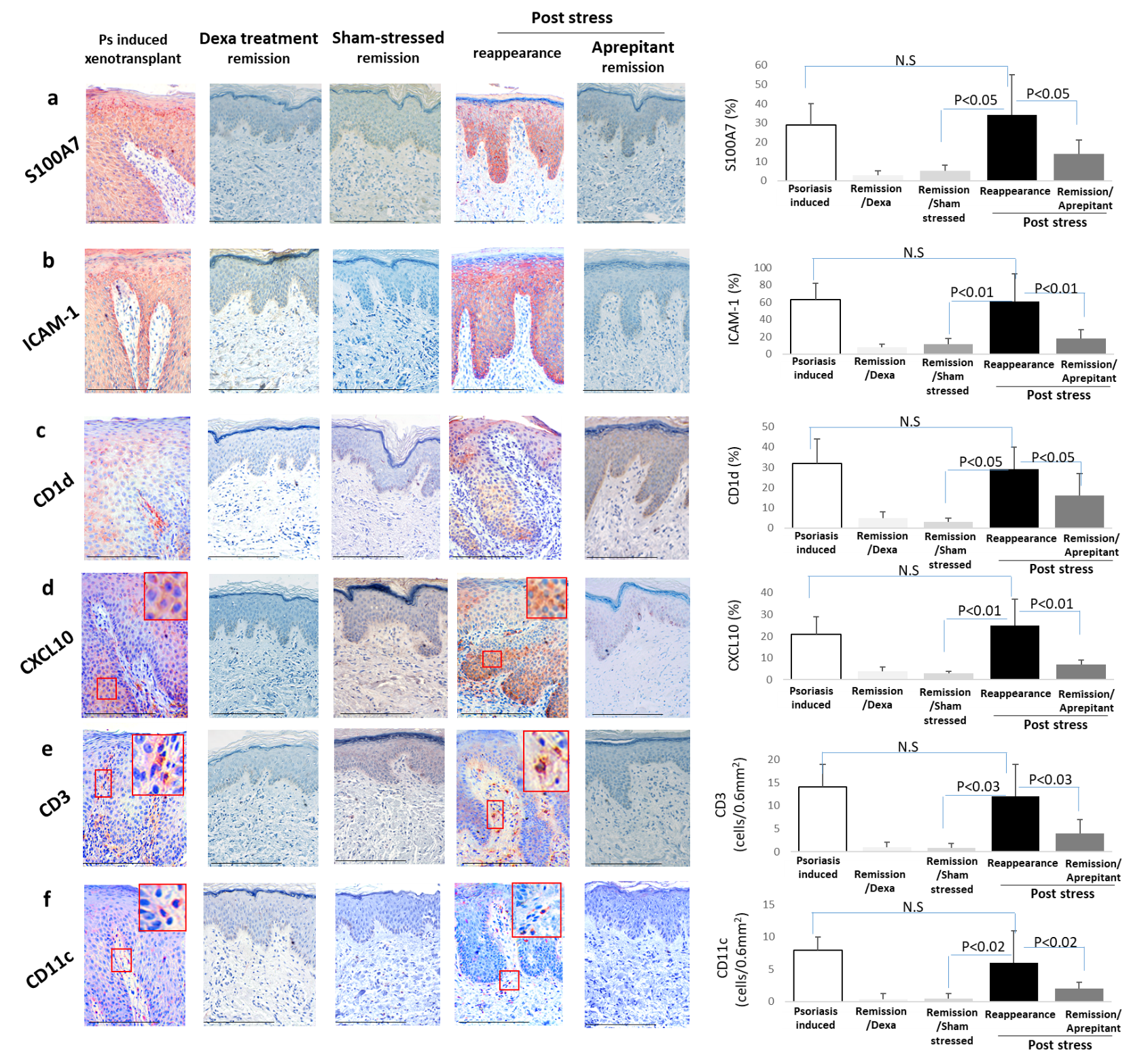
**Supplementary Figure 1**.

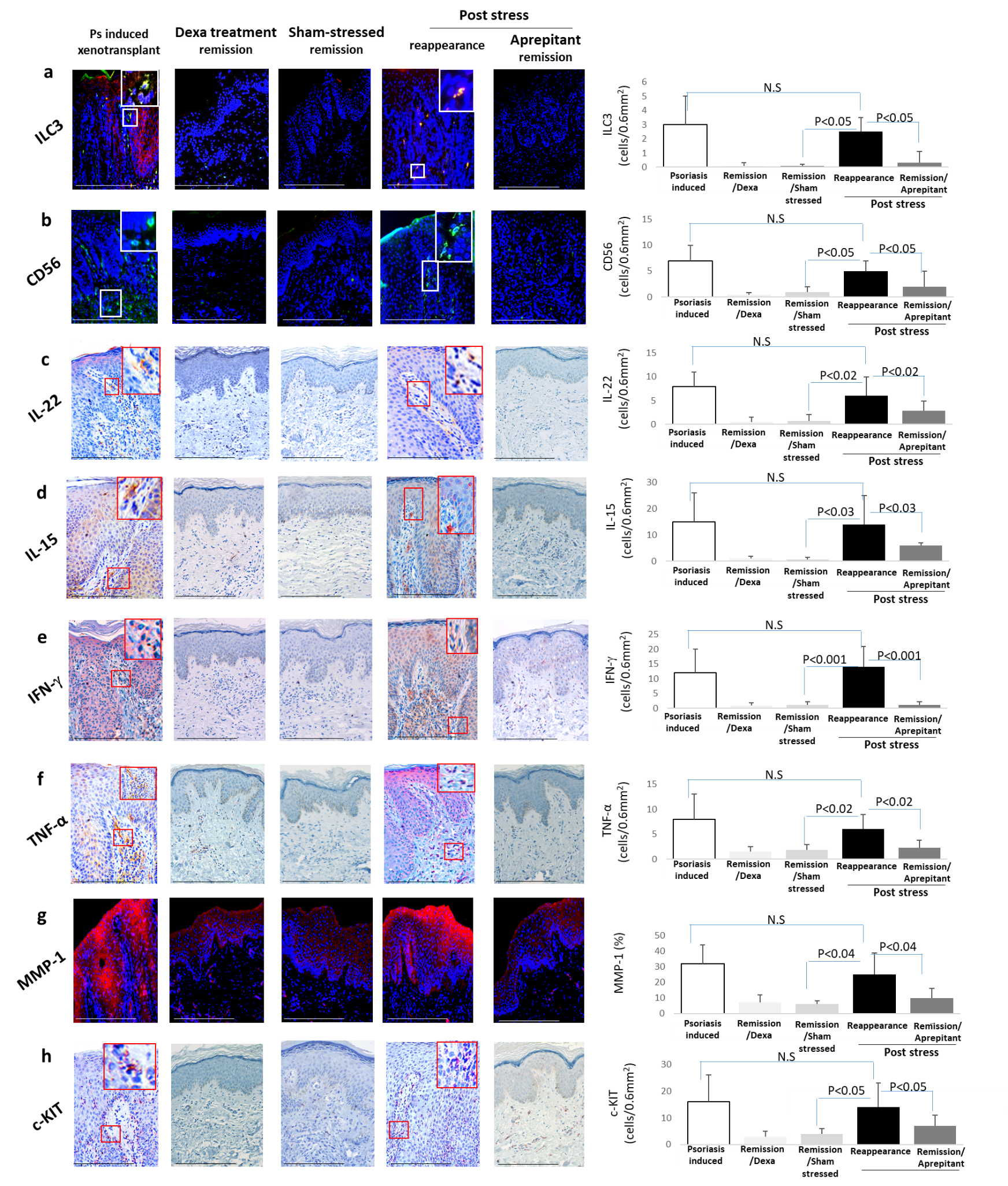
 **Supplementary Figure 2.**

**Supplementary Table 1. Dynamics of Psoriatic Marker Expression Across Different Treatment and Stress Conditions**

|  | **Group**  **Marker** | **Psoriasis induced xenotransplants** | **Psoriasis remission/**  **Dexa treatment** | **Psoriasis remission/**  **Sham-stressed** | **Psoriasis reappearance/**  **Post stress** | **Psoriasis remission/**  **Aprepitant treatment (post stress)** | **P value** |
| --- | --- | --- | --- | --- | --- | --- | --- |
| **Markers of Epidermal Proliferation (%)** | Epidermal Thickness (µm) | 573±123 | 231±32 | 254±76 | 515±160 | 320±110 | <0.02 |
|  | Ki-67 | 43±15 | 7±3 | 6±3 | 36±16 | 10±3 | <0.03 |
|  | K16 | 64±21 | 6±4 | 8±4 | 54±14 | 14±7 | <0.02 |
| **Anti-Microbial Protein (%)** | hBD2 | 44±18 | 12±7 | 10±4 | 34±13 | 19±12 | <0.05 |
|  | S100A7 | 29±11 | 3±2 | 5±3 | 33±21 | 14±7 | <0.05 |
| **Epidermal Immune Response Markers** | HLA-DR (%) | 65±24 | 13±7 | 10±6 | 58±21 | 16±10 | <0.04 |
|  | ICAM-1 (%) | 63±11 | 8±3 | 11±7 | 61±32 | 18±11 | <0.01 |
|  | CD1d (%) | 32±12 | 5±3 | 1±1 | 29±11 | 16±11 | <0.05 |
|  | CXCL10* | 15±6 | 2±0.5 | 2±1 | 17±5 | 3±2 | <0.01 |
| **Dermal Immune Cells Infiltrate*** | CD3 | 14±5 | 1±1 | 0.8±1 | 12±7 | 4±3 | <0.03 |
|  | CD8 | 16±12 | 1.4±1 | 1.2±1 | 14±8 | 3.4±2 | <0.03 |
|  | CD11c | 8±2 | 0.4±1 | 0.5±1 | 6±5 | 2±1 | <0.02 |
|  | CD56 | 7±3 | 0.4±0.5 | 1±1 | 5±2 | 2±3 | <0.05 |
|  | ILC3 | 3±2 | 0.1±0.2 | 0.1±0.1 | 2.5±1 | 0.3±0.8 | <0.04 |
| **Cytokines Expressed by Dermal cells*** | IL-22 | 8±3 | 0.5±12 | 0.7±1.3 | 6±4 | 2.9±2 | <0.02 |
|  | IL-17 | 13±8 | 1.2±0.9 | 1±0.5 | 10±6 | 5±2 | <0.05 |
|  | IFN-γ | 12±8 | 1±1 | 1.2±1 | 14±7 | 1±1 | <0.001 |
|  | TNF-α | 8±5 | 1.5±1 | 2±1 | 6±3 | 2.3±1.5 | <0.02 |
|  | IL-15 | 15±11 | 1±1 | 0.6±1 | 14±11 | 6±1 | <0.03 |
| **Mast cells*** | c-KIT | 16±10 | 3.5±1.4 | 4±2 | 14±9 | 7±3.2 | <0.05 |
|  | Tryptase | 15±12 | 3±2 | 2±2 | 18±14 | 6±4 | <0.02 |
| **Angiogenic Factor** | VEGF-A (%) | 43±23 | 10±6 | 12±4 | 41±28 | 21±11 | <0.05 |
| **Matrix Metalloproteinase** | MMP-1 (%) | 32±12 | 7±5 | 6±2 | 25±14 | 10±6 | <0.05 |
| **Neuro-regulatory Factors** | NGF (%) | 37±16 | 10±3 | 8±5 | 52±21 | 21±14 | <0.02 |
|  | NK-1R* | 15±6 | 31±2 | 5±3 | 18±7 | 9±3 | <0.04 |
|  | Substance P (%) | 24±215 | 1±0.5 | 2±1 | 29±11 | 12±4 | <0.04 |

* Number of cells/0.66mm^2^

Data are presented as the mean ± standard error of mean (SEM); p values of <0.05 were regarded as significant
